# Mechanisms of Chromosomal Instability in High-grade Serous Ovarian Carcinoma

**DOI:** 10.1101/727537

**Authors:** N. Tamura, N. Shaikh, D. Muliaditan, J. McGuinness, D. Moralli, M. A. Durin, C. M. Green, D. Bowtell, F. R. Balkwill, K. Curtius, S. E. McClelland

## Abstract

Chromosomal instability (CIN), the continual gain and loss of chromosomes or parts of chromosomes, occurs in the majority of cancers and confers poor prognosis. Mechanisms driving CIN remain unknown in most cancer types due to a scarcity of functional studies. High-grade serous ovarian carcinoma (HGSC), the most common subtype of ovarian cancer, is the major cause of death due to gynaecological malignancy in the Western world with chemotherapy resistance developing in almost all patients. HGSC exhibits high rates of chromosome aberrations and knowledge of causative mechanisms is likely to represent an important step towards combating the poor prognosis of this disease. However, very little is known about the nature of chromosomal instability exhibited by this cancer type in particular due to a historical lack of appropriate cell line models. Here we perform the first in-depth functional characterisation of mechanisms driving CIN in HGSC by analysing eight cell lines that accurately recapitulate HGSC genetics as defined by recent studies. We show, using a range of established functional CIN assays combined with live cell imaging and single molecule DNA fibre analysis, that multiple mechanisms co-exist to drive CIN in HGSC. These include supernumerary centrosomes, elevated microtubule dynamics and DNA replication stress. By contrast, the spindle assembly checkpoint was intact. These findings are relevant for developing therapeutic approaches to manipulating CIN in ovarian cancer, and suggests that such approaches may need to be multimodal to combat multiple co-existing CIN drivers.

## Introduction

The vast majority of solid tumours exhibit chromosomal instability (CIN), the continual gain and loss of chromosomes or parts of chromosomes^1,2^. CIN can drive tumour heterogeneity and clonal evolution, and is thought to contribute to chemotherapy resistance in many cancer types including ovarian cancer^3–5^. Knowledge of the defective cellular pathways that underlie CIN would enable strategies to target cancer cells using synthetic lethal or CIN-limiting approaches^6^, in addition to providing new diagnostic or prognostic tools. However, to date the causes of CIN in cancer remain ill-defined. It is thought that defective chromosome attachment to the mitotic spindle due to aberrant mitotic microtubule dynamics could contribute to CIN in cancer cell lines^7–10^, potentially driven by alterations in spindle protein abundances^7^ or genetic lesions in Aurora A, BRCA1 or Chk2^10^. Loss of retinoblastoma protein (pRB) leading to cohesion defects and CIN has also been demonstrated in a sarcoma cell line^11^. Studies classifying CIN mechanisms in representative, cancer specific cell panels are currently limited to colorectal cancer^9,12^. In this cancer type, both DNA replication stress, the slowing or stalling of DNA replication, and error-prone mitosis due to elevated microtubule assembly rates have been demonstrated to contribute to CIN^9,12^.

High-grade serous ovarian carcinoma (HGSC) represents an important clinical challenge; despite initial positive responses to first-line platinum therapy, most patients unfortunately relapse, leading to a poor overall survival for this disease^13^. HGSC genomes bear the scars of chromosomal instability as evidenced by highly aberrant genomic landscapes^4,14,15^ and evidence of ongoing CIN has been demonstrated in ascites-derived HGSC cells^16^. There has been extensive interest in interpreting the mutational signatures encoded within tumour and cancer cell line genomes over recent years^17,18^. Such studies have made important progress in inferring potential cancer mutational mechanisms at both single nucleotide variant and more recently, chromosome-scale aberrations, particularly in ovarian cancer^19–22^. As a result, a spectrum of potential mutagenic mechanisms has been inferred to operate in HGSC^20,22^. However, apart from a high prevalence of mutations in homologous recombination (HR) genes, and near ubiquitous *TP53* mutations^14,23^, genetic drivers of CIN in this disease remain to be definitively elucidated. Additional possible genetic lesions that may contribute to CIN in HGSC are Aurora A amplification and Cyclin E *(CCNE1)* amplification^24^. *RB1* mutations and inactivating gene breakage events are also present in 17.5% HGSC tumours^15^. Importantly, it has also been shown that BRCA-mutated tumours can acquire HR reactivating mutations^25,26^ highlighting the need for functional analysis in defining ongoing CIN mechanisms.

To date there has been a lack of functional characterisation using appropriate cell line models for HGSC. Recent advances have been made in defining the many and distinct subtypes of ovarian cancer, including genomic approaches allowing the classification of available tumour-derived cell lines into suitable models^27–29^. We therefore undertook the first systematic and comprehensive functional characterisation of a curated panel of HGSC cell lines to define mechanisms driving chromosomal instability. We demonstrate that all cell lines exhibit extensive, ongoing CIN in the form of high rates of numerical and structural chromosome defects, and chromosome segregation errors. We find gross defects in multiple pathways controlling chromosome stability, including supernumerary centrosomes, elevated microtubule dynamics, and replication stress. We show that either suppressing MT dynamics using low doses of taxol, or limiting replication stress using nucleoside supplementation, reduce chromosome segregation errors and CIN. This new knowledge regarding mechanisms driving CIN in HGSC provides a first step to designing new approaches to treat high grade serous ovarian carcinoma, including for example determining whether limiting CIN could prevent CIN-driven chemotherapy in HGSC^6^.

## Results

### Numerical and structural chromosome defects, and persistent chromosome segregation errors in HGSC cell lines

Cancer-derived cell lines have proven a useful resource to measure ongoing mechanisms driving chromosomal instability^7,9,12,30^. We assembled an array of eight HGSC lines, five previously identified as suitable models for HGSC (Cov318, Kuramochi, Ovkate, Ovsaho, Snu119)^27^ and three obtained from recent confirmed HGSC patients (AOCS1^31^, G33^31^ and G164) (**Table S1**). As tissue-type specific controls we obtained two h-TERT-immortalised fallopian tube serous epithelial cell lines (FNE1 and FNE2^32^), representing the likely tissue of origin of HGSC^33^. We performed whole genome sequencing (Methods) to characterize the extent of structural abnormality in the HGSC lines. First, to visualise copy number gains and losses, we performed copy number segmentation^34^ and computed the DNA copy number profiles (**Figure 1a**; see Methods for details). Since matching normal DNA for each line was not available, we down-sampled our sequencing data to 0.1x^35^ and utilised the R Bioconductor ACE package to estimate absolute DNA copy numbers. This analysis also computed the most likely models of ploidy (**Figure S1a**), and we further confirmed these using chromosome counting from metaphase chromosome spreads (**Figure 1b,c: Figure S1b**) and FACS ploidy analyses (**Figure S1c**). Similar to HGSC genomes available in the TGCA dataset^14,27^, the HGSC lines displayed complex copy number profiles (**Figure 1a**). There was a notable range in ploidy between cell lines, varying between near-diploid (2n) to near-tetraploid (4n) (**Figure 1c**). To examine chromosome alterations at a single cell level we performed multiplex-FISH (M-FISH) on metaphase chromosome spreads for FNE1 and Kuramochi cell lines. As expected, FNE1 were near diploid (**Figure 1d**; **Table S1**). By contrast, Kuramochi cells displayed a high prevalence of numerical and structural alterations that were highly heterogeneous between individual cells (**Figure 1d** and **Figure S1d**). Metaphase chromosome spreads analysed with centromeric fluorescence *in-situ* hybridisation (FISH) probes also demonstrated the presence of structural chromosome aberrations (dicentric and acentric chromosomes) in most HGSC cell lines (**Figure S1e,f**). Our evaluation of the inter-centromere distance, which provides a measure of sister chromatid cohesion defects^36^, suggested chromosome cohesion was normal in all HGSC lines with the possible exception of COV318 (**Figure S1g**).

**Figure 1:**
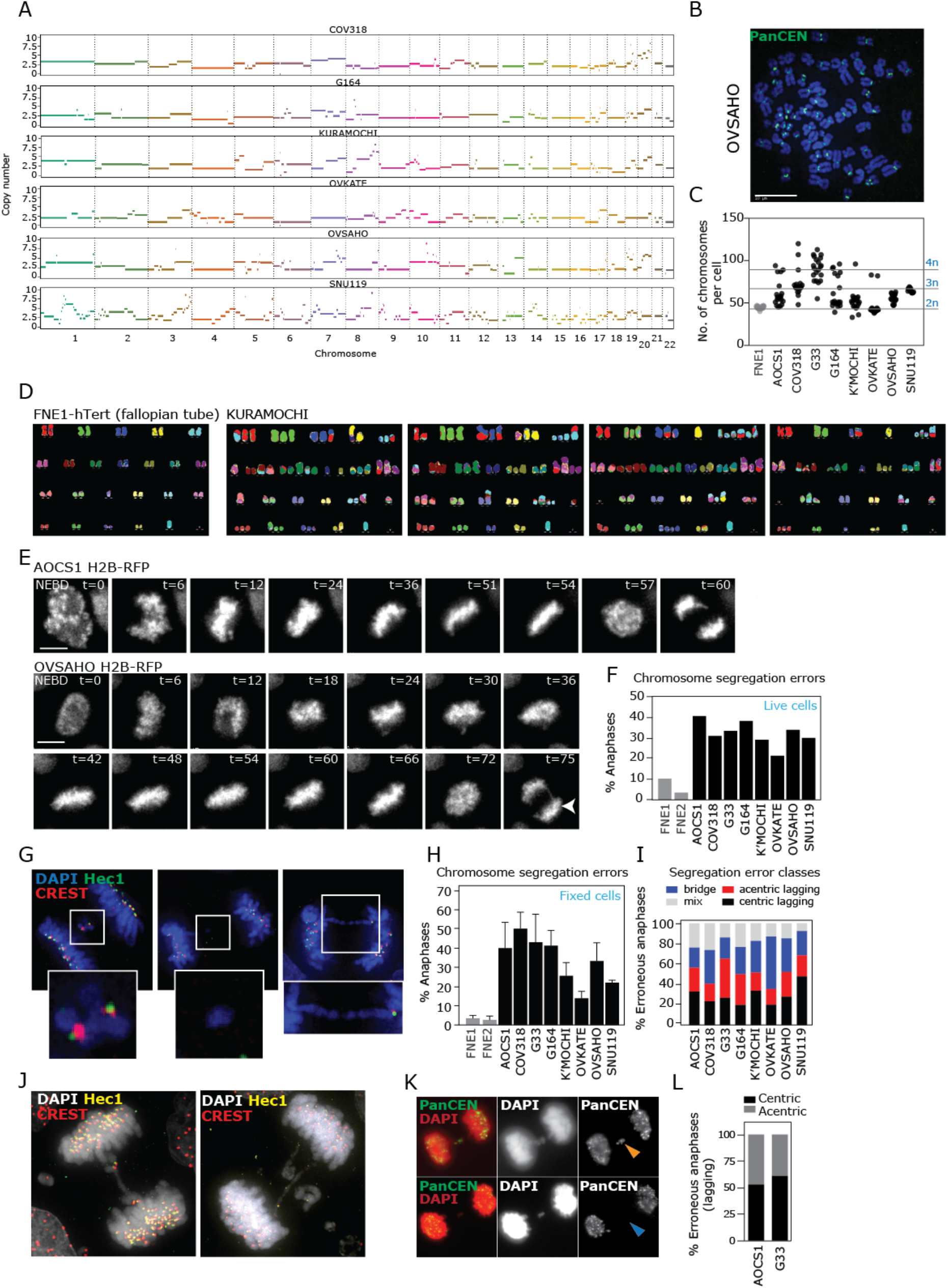
HGSC cell lines display numerical and structural chromosome defects and persistent chromosome segregation errors. **A)** DNA copy number profiles computed from WGS data for six HGSC cell lines. **B)** Representative image of metaphase spread from Ovsaho. Pan-centromere FISH staining indicated in green (panCEN). **C)** Ploidy of each cell line was derived from metaphase spread analysis. For each cell line, twenty metaphase spreads were analysed across two experiments. **D)** M-FISH analysis of metaphase spreads showing structural and numerical aberrations, with one example from FNE1 and four examples from Kuramochi. **E)** Stills taken from movies of two cell lines stably expressing H2B-RFP (white). Time in minutes is indicated post nuclear envelope breakdown (NEBD). **F)** Chromosome segregation error rates from live cell movies. Data shown is from two experiments. **G)** Immunofluorescence images of anaphase cells probed with antibodies to CREST (red, marks centromere) and Hec1 (green, marks kinetochore) exhibiting different classes of segregation errors; lagging centric, lagging acentric, anaphase bridge. **H)** Analysis of segregation errors of all cell lines from immunofluorescence images, from four to seven experiments (179-300 anaphase cells per cell line). **I)** Anaphase cells with errors from H were classified according to error type (either centric, acentric or bridge errors or a mixture of any or all three types). **J)** Example Cov318 anaphase cells with multiple errors per cell. **K,L)** FISH with centromeric probes (PanCEN; green) was used to define whether lagging chromatin (red) was positive (centric) or negative (acentric) for DNA centromere sequence, quantified in (L) for 104 (Aocs1) and 108 (G33) anaphases. All scale bars 10 μm.

*TP53* mutations occur in 96% of HGSC tumours^14^. Accordingly, all six sequenced HGSC cell lines exhibited *TP53* mutations and aberrant p53 protein expression compared to FNE1 cells (**Figure S1h**). We also noted heterogeneity between the cell lines in terms of common genomic features of HGSC, including *CCNE1* aberrations at the genetic and protein level (**Figure S1i,j**). We verified the known *BRCA2* mutation in Kuramochi^27^, but did not identify any other *BRCA1/2* mutations of known pathogenicity (**Table S2**). Some lines exhibited *BRCA1* or *BRCA2* copy number alteration (**Figure S1i**) although it is not clear whether this is sufficient to impair function. Our HGSC cell line panel thus recapitulates key genomic features of HGSC tumours and cell lines, and encompasses a range of ploidy, genetic and genomic alterations.

The diversity of chromosome alterations between individual Kuramochi cells, and the prevalence of structural and numerical chromosome alterations in all HGSC lines, suggested that chromosomal instability was ongoing. To directly test this, we created cell lines stably expressing mRFP-tagged Histone2B and monitored chromosome segregation during mitosis using live cell imaging. We observed the frequent occurrence of chromosome mis-segregation events in all HGSC cell lines (**Figure 1e,f**). Chromosome segregation errors were further examined using high resolution imaging of fixed cells to gain insights about the nature of mis-segregating chromatin (**Figure g-i**). We were struck by the very high frequency of chromosome mis-segregation in some lines, for example Cov318 exhibited errors in 50% of cells, with each cell typically displaying multiple errors often of different types (**Figure 1j**). Lagging chromatin from anaphase cells was sometimes negative for CREST-reactive serum (that marks centromeric proteins) and Hec1 kinetochore proteins, suggesting that some mis-segregation events were precipitated by structural chromosome alterations (**Figure 1g,i,j**). To verify the acentric nature of lagging chromatin we performed FISH using all-centromere-targeted probes in two cell lines which confirmed the presence of mis-segregating chromatin devoid of centromeric DNA sequence (**Figure 1k,l**). HGSC cell lines thus exhibit continual mis-segregation of both intact and structurally abnormal chromosomes, contributing to their high rates of numerical and structural CIN.

### HGSC cell lines prolong mitosis due to slow alignment of chromosomes to the metaphase plate

To test whether other aspects of mitosis were perturbed in HGSC we analysed mitotic progression kinetics using live cell imaging of H2B-RFP stable cell lines. Slight congression delays have been reported in colorectal cancer^9^ but it was hitherto unknown whether this feature occurs in other cancer types. Five HGSC cell lines exhibited significant delays in mitosis, as measured by the time from nuclear envelope breakdown to anaphase onset (**Figure 2a, b**). To test whether this was a consequence of faulty chromosome alignment to the metaphase plate, or a delay in completing satisfaction of the mitotic checkpoint, we scored the time at which all chromosomes had congressed to form a complete metaphase plate. The cell lines with delayed mitotic timing were also significantly delayed in their chromosomal congression (**Figure 2c**) demonstrating a defect in this process, rather than a checkpoint defect. There was no obvious correlation between higher cell line ploidy and slowed congression, suggesting this phenomenon is unconnected with the number of chromosomes present. The fact that slow congression kinetics were accompanied by a delay in anaphase onset suggested that congression errors were capable of mounting a robust mitotic checkpoint response. In line with this, we observed loading of two mitotic checkpoint proteins, Mad2 and BubR1 on uncongressed chromosomes in all four cell lines tested (**Figure 2d,e**; **Figure S2a,b**). Taken together these data indicate that the spindle assembly checkpoint is intact in HGSC.

**Figure 2:**
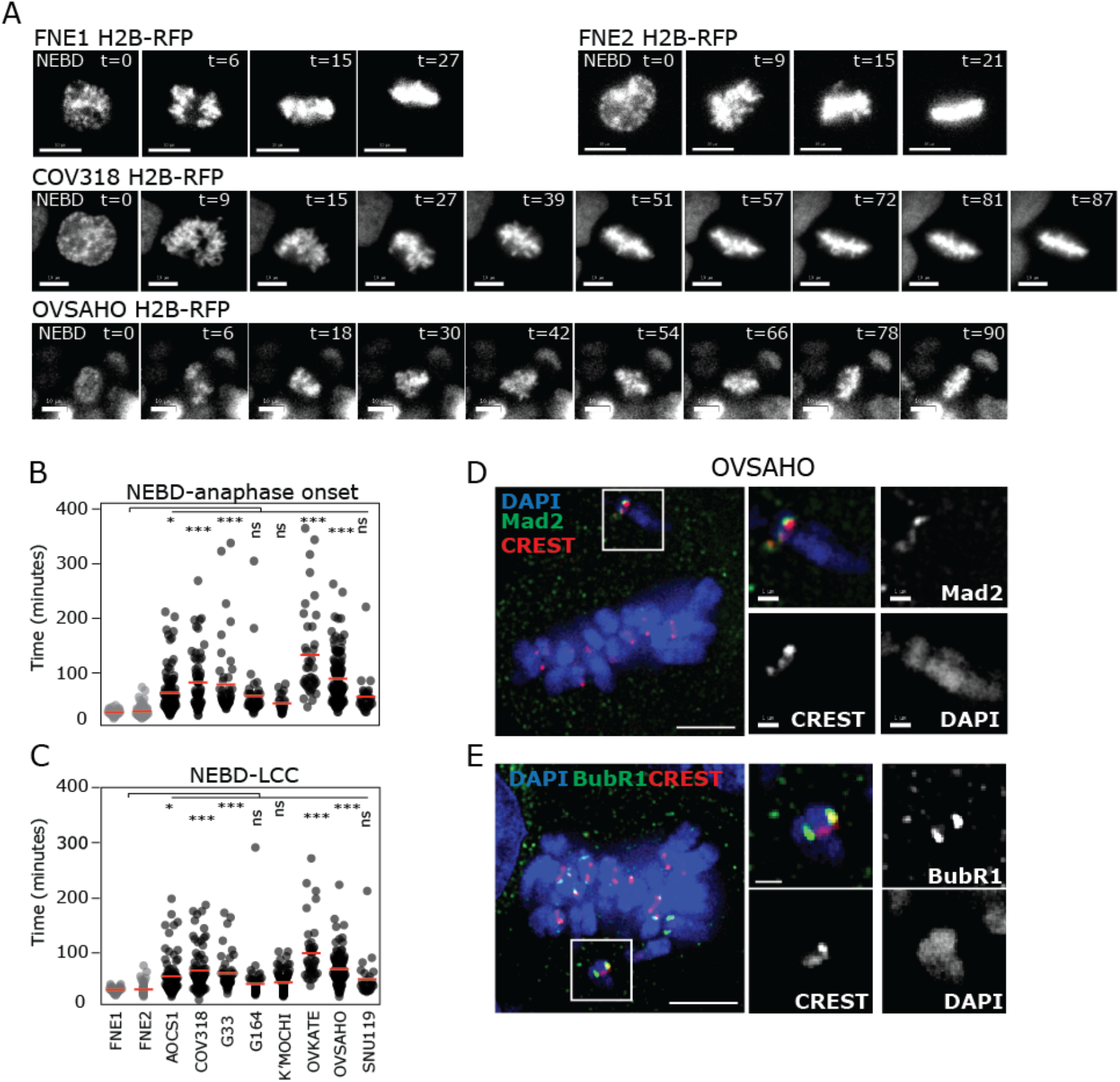
HGSC cell lines prolong mitosis due to slow alignment of chromosomes to the metaphase plate. **A)** Stills taken from movies of cell lines stably expressing H2B-RFP (white). Time in minutes is indicated post nuclear envelope breakdown (NEBD). **B)** Time taken for cells to progress from NEBD to anaphase onset. Each circle represents timing for one cell. Data taken from two independent experiments; 22-126 cells analysed per cell line. Statistical difference between HGSC cancer cell lines and FNE1 control is indicated. **C)** Time taken for cells to progress from NEBD to last chromosome congressed (LCC) to the metaphase plate. Data taken from two independent experiments. 26-109 cells analysed per cell line. Statistical test is one way Anova with correction for multiple testing. Differences between HGSC cancer cell lines compared to FNE1 are shown. **D,E)** Examples of immunofluorescence images of prometaphase cells with uncongressed chromosomes, stained for SAC components (Mad2 or BubR1; green) and centromeres (CREST; red). White boxes indicate uncongressed chromosome, with enlarged image to the right. Scale bars 5 μm or 1 μm (zooms). Asterisks denote significance (*p<0.05, ***p<0.001 or non-significant (ns)).

### Supernumerary centrosomes and aberrant MT dynamics contribute to chromosome segregation errors in HGSC

The significant delays in chromosome congression, and the presence of some apparently whole (centric), lagging chromosomes in anaphase (**Figure 1g,i,j**) suggested that the mitotic machinery was disrupted in HGSC. Centrosome abnormalities have been detected in ovarian cancer^30,37^, and can promote the formation of multipolar spindles^38^. Although multipolar spindles can resolve to a pseudo-bipolar spindle in a process known as centrosome clustering^39^, this can elevate the frequency of incorrect chromosome attachments to the spindle and increase chromosome segregation errors^38,40^. We therefore quantified centrosome and spindle defects from mitotic cells using immunofluorescence and antibodies against centrioles (centrosome cores; two per centrosome). A majority of cell lines exhibited supernumerary (>4) centrioles per cell (**Figure 3a,b**). Cell lines with extra centrosomes accordingly exhibited defects in mitotic spindle polarity (**Figure 3c,d**). To investigate whether centrosome amplification was associated with elevated chromosome segregation errors in HGSC we compared error rates between anaphase cells with 4, or more than 4 centrioles. In two lines (G33 and G164), the presence of extra centrosomes correlated with a higher rate of chromosome segregation errors (**Figure 3e,f**). By contrast, no association was observed in AOCS1 cells between extra centrosomes and elevated segregation errors, suggesting that extra centrosomes might contribute to elevated chromosome mis-segregation in some but not all cell lines.

**Figure 3:**
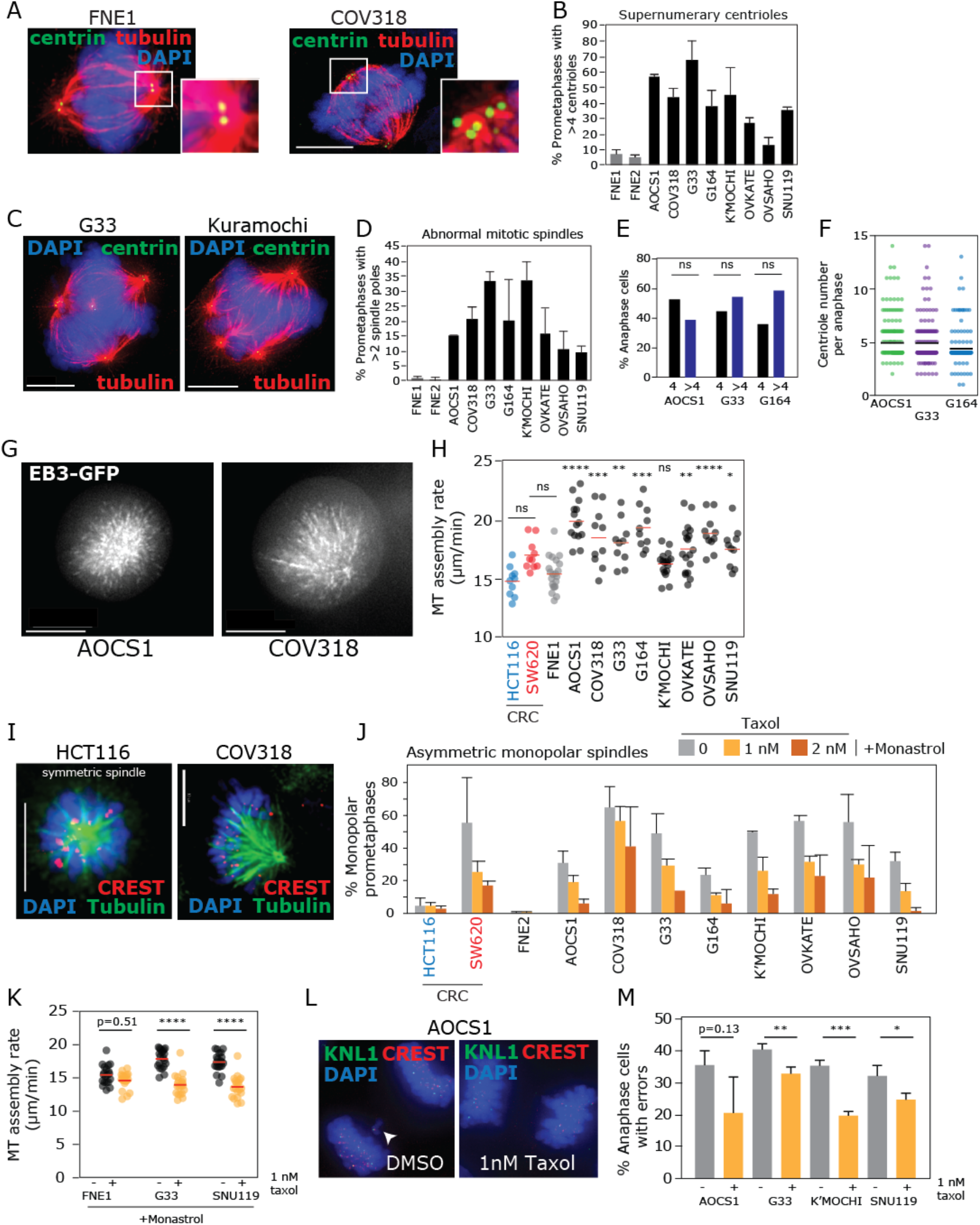
Supernumerary centrosomes and aberrant MT dynamics contribute to HGSC CIN. **A)** Examples of immunofluorescence images of cells in prometaphase, probed with antibodies against centrin (green) to mark centrioles and tubulin (red) to mark microtubules. **B)** Quantification of percentage of prometaphase cells with abnormal (>4) centrioles. **C)** Examples of immunofluorescence images of prometaphase cells with abnormal mitotic spindles. **D)** Quantification of percentage of cells with abnormal mitotic spindles. **E)** Comparison of chromosome segregation error rates in anaphase cells with normal (4) and abnormal (>4) centrioles. T-test between pairs is shown. **F)** Quantification of centriole number in anaphase cells from three cell lines. Data in E and F are summed from two independent experiments (152-213 anaphases per cell line in total). **G)** Stills taken from live movies of cells expressing EB3-GFP, arrested in mitosis using monastrol. **H)** Analysis of microtubule dynamics from EB3-GFP live imaging tip tracking assay. Statistical test is one way Anova with correction for multiple testing. Differences between HGSC cancer cell lines compared to FNE1 are shown. **I)** Examples of immunofluorescence images of prometaphase cells arrested with monastrol, showing normal (HCT116) or abnormal (Cov318) spindle morphology. **J)** Quantification of abnormal spindle morphology in cells arrested with monastrol in conjunction with 0, 1 or 2 nM taxol. **K)** Measurements of microtubule dynamic rates in cells expressing EB3-GFP, arrested with monastrol and treated with either DMSO or 1 nM taxol. **L,M)** Analysis of segregation error rates of cells after 2 hr treatment with DMSO or 1 nM taxol. T-tests were performed in K) and M) between pairs as indicated. Scale bars 10 μm (1 μm in zooms). Asterisks denote significance (*p<0.05, **p<0.01, ***p<0.001 ****p<0.0001, or non-significant (ns). P values close to 0.05 are indicated numerically.

Elevated microtubule (MT) dynamics leading to delayed chromosome congression, chromosome segregation errors and CIN was recently described in colorectal cancer as a result of either Aurora A overexpression^9^ or the deregulation of the Chk2-BRCA1 axis^10,41^. Since HGSC also carries potential Aurora A defects due to a common amplification of the chromosomal region carrying the Aurora A gene (*AURKA*, 20q^27^; **Figure 1a**; **Figure S3a**) and given the known *BRCA1/2* mutations frequently observed in HGSC, we tested whether MT dynamics were similarly altered in HGSC cell lines. To do this we transiently expressed the MT tip-tracking protein EB3 tagged with GFP using lentiviral transfection and filmed MT dynamics. To allow accurate quantification of mitotic spindle MT dynamics, cells were treated with the Eg5 inhibitor Monastrol to generate monopolar spindles as previously described^9^ (Figure 3g and Movie S1). We included the colorectal cancer cell lines SW620 (CIN-positive) and HCT116 (CIN-negative) as controls^9^. MT assembly rates were significantly elevated in all HGSC cell lines (except Kuramochi) compared to FNE1 cells (Figure 3h) with most lines notably displaying MT assembly rates well above CIN-positive SW620 colorectal cancer cells. To test whether elevated MT dynamics was causative for CIN in HGSC we treated cells with a low dose of the MT stabilising agent, taxol, previously demonstrated to suppress MT assembly rates in colorectal cancer^9^. To first establish whether low dose taxol could suppress elevated MT assembly rates in HGSC, we took advantage of an proxy read out: monopolar mitotic spindles frequently orient asymmetrically when MT assembly is elevated^42^ (**Figure 3i**). Using this assay, we confirmed that low dose taxol treatment reduced MT assembly rates in HGSC cell lines (**Figure 3i, j**). We also directly confirmed the reduction in MT assembly rates using the EB3-GFP tip tracking assay for two HGSC lines (**Figure 3k**). Importantly, restoration of MT assembly rates reduced chromosome segregation errors in four HGSC cell lines tested (**Figure 3l,m**) independent of any effect on proliferation (**Figure S3b**) demonstrating that aberrant MT dynamics contribute to chromosome mis-segregation in this HGSC cell line panel.

### HGSC cell lines exhibit replication stress that contributes to chromosome mis-segregation and CIN

The presence of acentric lagging chromatin structural chromosome defects described above and the known roles of homologous recombination proteins frequently mutated in HGSC including BRCA1 and BRCA2 in protecting against replication stress^43–46^ suggested that HGSC cell lines may experience replication stress. Replication stress, the slowing or stalling of DNA replication, is known to occur in multiple cancer types^47,48^ and was previously shown to contribute to CIN in colorectal cancer by generating unstable chromosome structures that are mis-segregated during mitosis in the form of acentric fragments and chromatin bridges^12^. To test whether replication stress occurs in HSGC lines we examined several known hallmarks of replication stress, namely; elevated prometaphase DNA damage, 53BP1 bodies in G1 cells and ultrafine anaphase bridges^12,49^. All eight HGSC cell lines exhibited replication stress compared to FNE1 cells, although to differing degrees and sometimes with a non-uniform affect across hallmarks of replication stress (**Figure 4a-f**). We also directly examined replication fork speed using single DNA fibre labelling. All cell lines exhibited reduced replication fork rates, with some cell lines (e.g. G164 and Ovkate) replicating DNA at almost half the rate of control FNE cells (**Figure 4g**; **Figure S4**). Genome content did not appear to affect replication speeds *per se,* since G33 cells (near tetraploid) exhibited the least reduced fork speed, and Ovkate cells (diploid) exhibited the slowest replication rate. To test whether elevated replication stress contributes to chromosome mis-segregation and CIN in HGSC we sought to reduce chromosome mis-segregation by reducing replication stress using nucleoside supplementation as previously described^12,50^. Four HGSC cell lines (Cov318, G164, Kuramochi and Ovsaho) responded to this treatment with small, but reproducible reductions in chromosome segregation errors (**Figure 4h,i**). Interestingly, AOCS1 and G33 tended towards an increased segregation error rate despite no detrimental effects on FNE cells (**Figure 4i**) possibly due to intrinsic differences between HGSC cell lines in their ability to balance cellular nucleotide levels. To test whether suppression of replication stress could translate into a reduction in karyotypic heterogeneity, we grew colonies from single cells and performed centromeric FISH to measure within-clone deviation in centromere number. Kuramochi and Ovsaho displayed reduced karyotypic heterogeneity following single cell colony derivation in the presence of nucleosides (**Figure 4j,k**) Cov318 did not show a reduction in colony mode deviation despite a reduction of chromosome segregation errors. Effects of nucleoside supplementation on chromosome segregation and karyotypic heterogeneity were independent of any effects on proliferation (**Figure S3b**). Taken together these data demonstrate the presence of replication stress that contributes, in at least a subset of lines, to chromosome segregation errors and CIN in HGSC.

**Figure 4:**
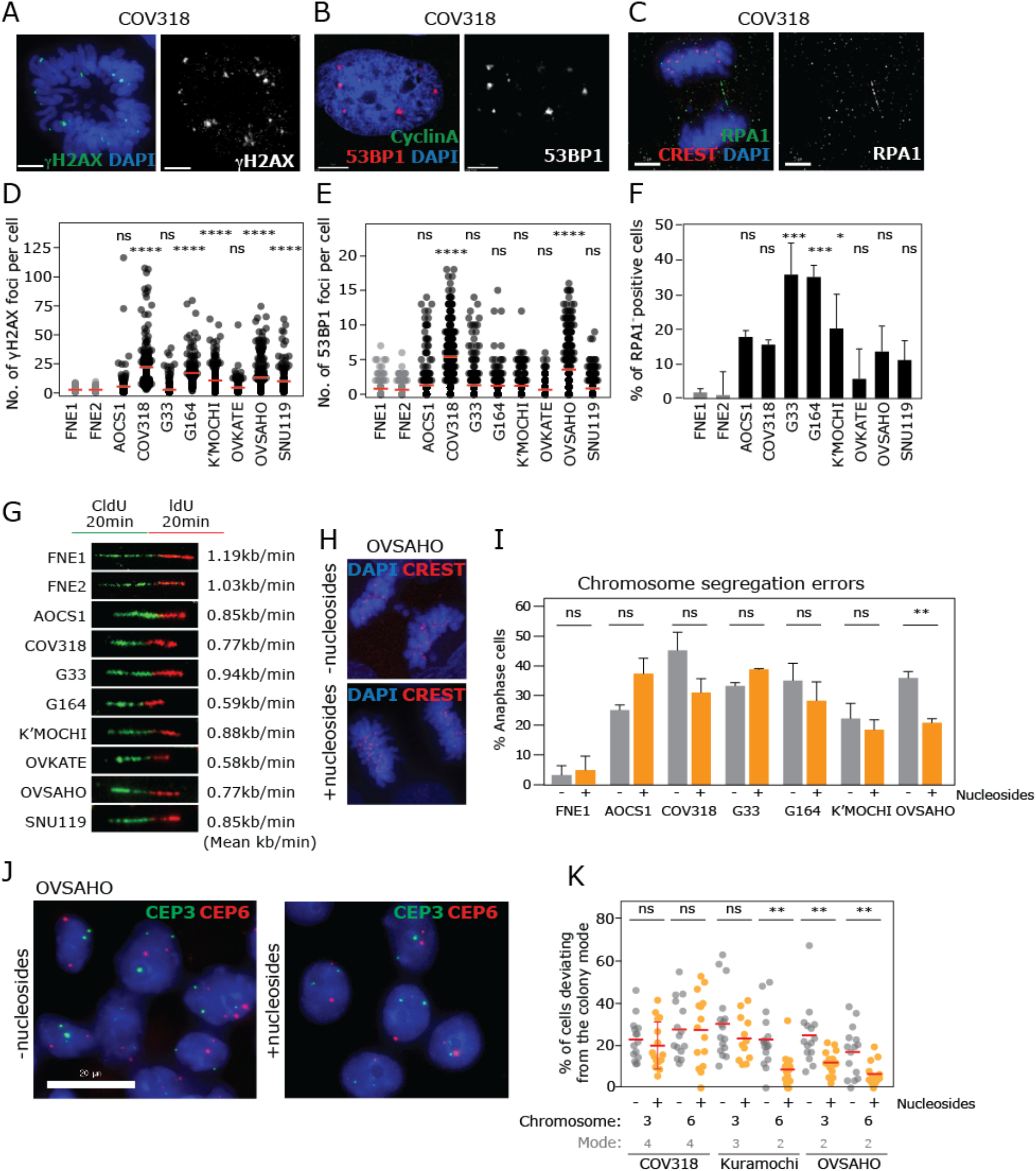
HGSC cell lines exhibit replication stress that contributes to chromosome mis-segregation and CIN. **A)** Example immunofluorescence (IF) image of prometaphase cell stained for γH2AX. **B)** Example of IF image of a G1 (Cyclin A negative) cell stained for 53BP1 bodies. **C)** Example IF image of an anaphase cell with an ultrafine bridge as demonstrated by a DAPI-negative, RPA bridge. **D,E,F)** Quantification of percentage of cells demonstrating phenotypes from A, B and C respectively. Statistical tests are one way Anova with correction for multiple testing. Differences between HGSC cancer cell lines compared to FNE1 are shown. **G)** Replication fork rates as measured by DNA fibre assays. **H, I)** Segregation error rates of untreated cells vs. cells treated with nucleosides. T-test between pairs is shown in I). **J)** Examples of clonal FISH images, with cells stained for probes against centromere enumeration probes (CEP) of chromosomes 3 (CEP3; red) and 6 (CEP6; green). **K)** Analysis of numerical CIN using clonal FISH without or with nucleoside treatment. For each cell line, each circle indicates percentage of cells in a colony deviating from modal value (modes given below graph) of that chromosome in that colony. T-test between pairs is shown. Asterisks denote significance (*p<0.05, **p<0.01, ***p<0.001 ***p<0.0001, or non-significant (ns). P values close to 0.05 are indicated numerically.

## Discussion

Here we have performed the first systematic and comprehensive functional analysis of mechanisms driving chromosomal instability in a panel of representative HGSC cell lines. All eight lines demonstrate extensive ongoing CIN in the form of chromosome mis-segregation that is associated with multiple mechanisms, including elevated microtubule dynamics, centrosome amplification, and replication stress (see **Figure S5a** for a summary of phenotypes across all cell lines). A striking finding of our study is that compared to CRC cell lines, HGSC cell lines exhibit a greater number of co-operating CIN mechanisms, and each CIN mechanism often operates at more extreme levels (for example slower replication fork rates and higher MT assembly rates than CRC cell lines). Moreover multiple CIN mechanisms likely exist within single HGSC cells as evidenced by the presence of chromosome segregation errors of multiple types per cell (**Figure 1g-j**). This complexity explains the partial reductions in CIN obtained using CIN limiting experiments (low dose taxol and nucleoside supplementation), and will be important to consider when designing approaches to target CIN therapeutically in this disease.

The excessive and complex nature of HGSC CIN is in accordance with the high number of breakpoints observed in HGSC genomes compared to colorectal and pancreatic cancers (**Figure S5b**). Interestingly, despite much attention on homologous recombination defects and replication stress in HGSC, a key driver of CIN in this panel appeared to be defects in mitosis, namely elevated MT dynamics and supernumerary centrosomes. We also noted that HGSC lines were not entirely uniform in the extent to which each CIN mechanism was manifest, and moreover the precise nature of replication stress varied between lines (**Figure S5a**). Given the differences in genomic alterations between HGSC patients we suggest that CIN mechanisms and the balance between multiple CIN mechanisms may vary not only between cancer types, but also between HGSC patients.

What genetic alterations might indicate the mitotic and replicative defects that culminate in CIN in this cancer type? Oncogene induced replicative and mitotic stress can result from a myriad of known oncogenes and tumour suppressor genes^51,52^ however HGSC harbours relatively few recurrent mutations. Centrosome amplification has been observed in many cancer types, although its origin is still unclear. It is possible that whole genome doubling (WGD) events caused by cytokinesis failure could generate both centrosome amplification and increases in chromosome ploidy. Indeed WGD is estimated to occur particularly frequently in HGSC^53,54^. However, many cells exhibited centriole numbers exceeding eight (the expected consequence of cytokinesis failure) (**Figure 3f**) and some centrosome amplified lines were close to diploid (e.g. Kuramochi), suggesting potential alternative routes to centrosome amplification. One such cause could be replication stress itself, since slow replication can cause extra centrosomes in HR deficient cells^55^.

All HGSC lines except Kuramochi exhibited markedly elevated microtubule assembly rates. In CRC elevated MT assembly rates are proposed to occur as a result of overactive Aurora kinase A^10,41^ or a deregulated BRCA1-Chk2 axis^9,10^. None of our eight cell lines exhibited *BRCA1* point mutations, although some lines displayed evidence of partial *BRCA1/2* copy number loss (**Figure S1i**). All lines except Snu119 however, displayed copy number gain of the *AURKA* locus (**Figure S3a**), suggesting that in this panel at least, *AURKA* gene amplification may be a key driver of MT over-assembly, mitotic abnormalities and CIN in HGSC. It is also possible that other genetic lesions could generate elevated MT assembly rates, for example DNA-PKcs has also recently been identified as upstream of the BRCA1-Chk2 axis^56^. Supernumerary centrosomes could also contribute to elevated MT assembly rates as a result of increased MT nucleation^57^, suggesting the intriguing possibility that extra centrosomes might mediate chromosome mis-segregation via mechanisms over and above their canonical role in promoting abnormal geometry during spindle formation.

Alongside mitotic defects, all eight HGSC cell lines also exhibited replication stress, as evidenced from cell biological hallmarks of replication stress and/or single molecule DNA fibre analysis. The partial genomic loss of *BRCA1* or *BRCA2* (**Figure S1i**) could contribute to replication stress and CIN in this panel. Interestingly, we noted differences between cell lines in terms of their response to nucleoside supplementation. One possibility is that the nature of replication stress may vary depending on the spectrum of genetic lesions present. For example, *CCNE1* amplification has been associated with replication stress due to perturbation of nucleotide pools as a consequence of unscheduled entry into S phase^50^. Two cell lines (Ovsaho and Cov318) display *CCNE1* amplification at the genomic and protein levels, and display reduced segregation errors upon nucleoside supplementation. It is therefore possible that *CCNE1* amplification contributes to replication stress in these lines. Cyclin E protein is also upregulated across most of the cell lines compared to FNE1 in the absence of clear genomic amplification, a decoupling noted previously in HGSC^58^ (**Figure S1i**). There could therefore be an alternative explanation for the difference in response to nucleoside supplementation. For example, despite showing hallmarks of replication stress, Aocs1 and G33 were not rescued by nucleoside supplementation. It is possible that their replication stress is driven by a mechanism independent of imbalanced nucleoside levels, or that nucleotide metabolism in these lines is such that exogenous application rather causes an adverse effect. Indeed chromosome segregation error rates slightly increase upon nucleoside supplementation for these lines (**Figure 4i**).

A potential caveat of this study is that there may be differences in the behaviour of cell line models compared to cells within tumours in patients. To mitigate against this, we selected a panel of HGSC whose genetic and genomic features recapitulated HGSC tumours^27–29^. Moreover, the behaviour of this cell line panel was notably different to previously characterised colorectal cancer cell panels that did not display overt chromosome congression delays or supernumerary centrosomes^9,12^. Therefore these CIN mechanisms are likely to be *bona fide* HGSC CIN mechanisms, although these should be validated *in vivo* in future studies. The lack of matched normal patient DNA samples also made some genetic analyses difficult. Herein we used various approaches to circumvent this issue. However, future studies will benefit from the acquisition of patient-specific matched normal DNA samples to permit more extensive genetic and genomic analyses, and to provide improved ability to link genotype to CIN phenotypes. Despite these issues, tumour-derived cell lines still represent the best currently available model to functionally connect genetic lesions, genomic alterations and CIN mechanisms. This study provides new understanding of the nature of CIN in HGSC, is directly comparable to existing knowledge of CIN mechanisms in other cancer types, and moreover lays the groundwork for future studies to validate mechanisms driving HGSC CIN *in vivo.* Our findings also have the potential to facilitate future research into synergising with patient-specific CIN mechanisms as a therapeutic strategy.

## Author contributions

NT performed most experiments and analysis. NS performed experiments in Figures 1, 2, 4, S1 and S3 and JM contributed to MT assembly rate experiments. DMu performed bioinformatic analysis supervised by KC. DMo and MD performed and analysed M-FISH experiments supervised by CG. FB provided cell lines and intellectual input and advice throughout the study. DB provided Aocs1 and intellectual input. SM conceived the experimental design and supervised all experimental work and analysis. NT, NS and SM wrote and edited the manuscript.

## Acknowledgements

We would like to thank Professor C. Swanton for the kind gift of SW620 and HCT116 cell lines. We would also like to thank all members of the McClelland laboratory for useful discussions and reading of the manuscript. We thank the CRUK Flow Cytometry Core Service at Barts Cancer Institute (Core Award C16420/A18066).

## Grant support

NT and DMu were funded by a Wellbeing of Women project grant (RG2040). NS was funded by the Pancreatic Cancer Research Fund (PCRF). CMG, DMo and MAD were funded by Wellcome core award 090532/Z/09/Z.). FB was funded by European Research Council (ERC322566) and Cancer Research UK (A16354, A25714).

## Supplementary materials

(one supplemental movie, two supplemental tables, five supplemental figures)

## Materials and Methods

### Whole Genome Sequencing

Sample processing and whole genome sequencing were carried out by Edinburgh Genomics for six of the eight cell lines. Samples were processed using Illumina TruSeq Nano libraries since genomic DNA concentration were low (range 5.386 – 17.251 ng/μ1). Sequencing was performed with Illumina HiSeq X instruments to an average depth coverage of 30X. Due to the nature of our cell line samples, matched normal tissue/blood samples were not available for analysis.

### WGS data pre-processing

FastQC was used to perform quality control (QC) of the raw .fastq files^59^. Each file was then aligned to the GRCh37 Human Genome Reference using BWA 0.7.8^60^. Resulting .sam files were compressed and converted to .bam files using samtools 1.9 ^61^. Picard version 2.6.0-SNAPSHOT was used to sort the .bam files and to mark duplicate reads (http://broadinstitute.github.io/picard/). Duplicate reads were removed and .bam files were indexed using samtools and picard respectively. Qualimapbamqc was used to conduct QC of the processed .bam files^62^. The GATK DepthOfCoverage and Picard CollectWgsMetrics tools were used to estimate genome wide coverage statistics^63^.

### Downsampling and absolute copy number estimation and visualization

To generate absolute copy numbers using unmatched tumour data, we first downsampled our 30x .bam files to 0.1x using samtools 1.9 and repeated the pre-processing as described above. Using these downsampled .bam files as input, we performed absolute copy number estimation using the R package ACE (15). Default binsizes were used (100kb, 500kb and 1000kb), with ploidy estimates of 2N, 3N and 4N. The loopsquaremodel function of ACE and resulting matrixplots were used to find the best ploidy and cellularity models. The ggplot2 package in R was used to plot the segmented copy number profiles (segmentation data from the resulting 100kb bins rds file was used).

### Gene-specific copy number alterations

Since matched normal data were not available for analysis, we calculated gene specific copy number changes using the formula mean gene coverage/mean genome coverage. Mean gene coverage was calculated using the bamstats05 tool and a custom bed file downloaded from the UCSC Table Browser^64,65^. A gene is indicated as having copy number gain if the mean gene coverage/mean genome coverage ratio is higher than 1.3 and having copy number loss if mean gene coverage/mean genome coverage < 0.7. We subsequently calculated this ratio using our data from ACE (mean segment copy number value of the segment containing that gene/ploidy) and found similar results (data not shown).

### Cell Lines

All HGSC and fallopian cell lines were sourced as detailed in **Table S1** and maintained at 37°C and 5% CO2. HCT116, SW620 (kind gift from C. Swanton) and Cov318 were maintained in DMEM High Glucose (Sigma); Kuramochi, G33, Ovkate, Ovsaho and Snu119 were maintained in RPMI (Sigma); both of these media types were supplemented with 10% FBS and 100 U Penicillin/Streptomycin. G164 cells were grown in DMEM F12 (Sigma) supplemented with 5% human serum (H4522, Sigma) and 100 U Penicillin/Streptomycin. FNE1/FNE2 cell lines were maintained in FOMI media (University of Miami) supplemented with cholera toxin (C8052, Sigma). H2B-RFP stable cell lines were generated after transfection with lentiviral construct H2B-RFP (Addgene 26001) and sorting for RFP-positive cells using flow cytometry. Cells were routinely tested for mycoplasma using MycoAlert PLUS Mycoplasma Detection Kit (LT07-710, Lonza) and visual inspection using DAPI staining at the microscope. Cells were passaged for a maximum of 8-12 weeks (approx. 10-14 passages).

### Proliferation assays

Equal numbers of cells were seeded into 96-well dishes. The next day, extra media was added (supplemented with either Embryomax nucleosides for final concentration of 10x, or paclitaxel for final concentration of 1 nM). Plate was then imaged over the course of one week using an IncuCyte® live cell analysis system. For each cell line, under each condition, the IncuCyte® calculated the percentage of confluency. We then reported the fold-change in confluency as a growth ratio (final confluency / starting confluency for that cell line grown under that condition).

### Metaphase spreads

Cells were arrested in colcemid for two hours, collected then resuspended in hypotonic solution (0.2% KCl, 0.2% Sodium Citrate) for 7 min at 37°C. Cells were pelleted and re-suspended in freshly-prepared 3:1 methanol-glacial acetic acid, then dropped onto slides.

### Clonal FISH

Cells were seeded onto slides at low density to ensure growth of colonies from single cells. Colonies were grown in presence or absence of nucleosides for four weeks then fixed for FISH. Cells in each colony were imaged and scored for centromere number, and percentage cells deviating from modal value for centromere of that colony was calculated.

### Small molecule inhibitors

100x Embryomax nucleosides (ES-008-D, Merck Millipore) were diluted in media to 10x final concentration. Taxol (Paclitaxel, P045, Cambridge Bioscience) was dissolved in DMSO and used at 1nM final concentration. Monastrol (Sigma) was dissolved in DMSO and used at 100μM.

### Immunofluorescence

Cells grown on coverslips were fixed with PTEMF (0.2% Triton X-100, 0.02 M PIPES (pH 6.8), 0.01 M EGTA, 1 mM MgCl_2_, 4% formaldehyde). After blocking with 3% BSA, cells were incubated with primary antibodies according to suppliers’ instructions. Antibodies were obtained from Abcam (Beta-tubulin (ab6046), CenpA (ab13939), Centrin 3 (ab54531), Cyclin A2 (ab16726), Hec1 (ab3613), RPA (ab79398)), Antibodies Incorporated (CREST (15-234-0001)), Bethyl Lab (Mad2 (A300 −300A)), Millipore (H2aX (05-636)), Santa Cruz (53BP1 (sc-22760)). Secondary antibodies used were goat anti-mouse AlexaFluor 488 (A11017, Invitrogen), goat anti-rabbit AF594, AF488 (A11012, A11008, Invitrogen), and goat anti-human AF647 (109-606-088-JIR, Stratech or A21445, Invitrogen). DNA was stained with DAPI (Roche) and coverslips mounted in Vectashield (Vector H-1000, Vector Laboratories).

### FISH

Fluorescent *In Situ* Hybridisation was carried out according to manufacturer’s instructions. In brief, cells on slides were fixed in 3:1 methanol:acetic acid, then put through an ethanol dehydration series (2 minutes in 70, 90, 100% ethanol) then air dried. Probe was added to slides which were heated to 72°C for 2 minutes, then left at 37°C overnight in a humid chamber. The next day, slides were washed in 0.25x SSC at 72°C for 2 minutes, then 2xSSC, 0.01% Tween at RT for 30sec. Slides were stained with DAPI then coverslips were mounted with Vectashield. Pan-centromere probe was purchased from Cambio (1695-F-02) and Centromere Enumeration Probes from Cytocell.

### M-FISH

Metaphase spreads from each cell line were hybridised with the M-FISH probe kit 24XCyte (Zeiss MetaSystems) following the manufacturer’s instructions. Briefly, the slides were incubated 30 minutes in 2xSSC buffer at 70°C, then allowed to cool at room temperature for 20 minutes. Following a 1 minute wash in 0.1XSSC, the cells were denatured in NaOH 0.07M for 1 minute, then washed in 0.1xSSC, and 2xSSC. The cells were dehydrated in an ethanol series, and air dried. The probe mix was denatured at 75 °C for 5 min, and preannealed at 37°C for 30 min. 6 μl of probe mix were applied to each slide, under a 18×18mm coverslip. The slides were incubated for 3 days at 37°C, then washed for 2 minutes in 0.4xSSC, at 72°C, and 30 second in 2xSSC, 0.05% Tween20, at room temperature, and finally mounted in DAPI/Vectashield (VectorLabs). Images were acquired on an Olympus BX-51 microscope for epifluorescence equipped with a JAI CVM4+ progressive-scan CCD camera, and analysed using the Leica Cytovision Genus v7.1 software (Leica). A minimum of 25 metaphases were karyotyped for each cell line.

### Fibre Assay

Fibres were prepared as described^66^. In brief, cells were pulse labelled with 25 μM CldU and 250μM IdU (Sigma) for 20 min. Cells were harvested and then lysed using 0.5% SDS, 20mM Tris-HCl pH 7.4, 50mM EDTA. Fibres were spread on slides and DNA detected using rat anti-BrdU and CldU, with secondary antibodies as above.

### Microscopy

Images were acquired using an Olympus DeltaVision RT microscope (Applied Precision, LLC) equipped with a Coolsnap HQ camera. Three-dimensional image stacks were acquired in 0.2 μm steps, using Olympus 100× (1.4 numerical aperture), 60× or 40× UPlanSApo oil immersion objectives. Deconvolution of image stacks and quantitative measurements was performed with SoftWorx Explorer (Applied Precision, LLC). H2B-RFP-labelled cells were live imaged in 4 well imaging dish (Greiner Bio-one). 20 μm z-stacks (10 images) were acquired using an Olympus 40× 1.3 numerical aperture UPlanSApo oil immersion objective every 3 min for 8 h using a DeltaVision microscope in a temperature and CO2-controlled chamber. Analysis was performed using Softworx Explorer. Microtuble assembly assays (see below) were performed in part using an Eclipse Ti-E inverted microscope (Nikon) equipped with a CSU-X1 Zyla 4.2 camera (Ti-E, Zyla; Andor), including a Yokogawa Spinning Disk, a precision motorized stage, and Nikon Perfect Focus, all controlled by NIS-Elements Software (Nikon)).

### Microtubule dynamics assay

Microtubule dynamics were analysed as described previously^9^. Briefly, assembly rates were calculated by tracking EB3-GFP protein foci in living cells. Cells were seeded onto glass-bottom dishes and transduced with virus containing pEGFP_EB3 (gift from S. Godinho) Cells were treated with Eg5 (Kif11) inhibitor monastrol (67 μM, Sigma) for 2 hr. Cells were then imaged on the DeltaVision microscope or using an Eclipse Ti-E inverted microscope (Nikon). Cells were imaged every 2 s using four sections with a Z-optical spacing of 0.4 μm. Average assembly rates (micrometres per minute) were calculated using data for 20 individual microtubules per cell for 10-20 cells.

### Western blotting

Cell lysates were prepared using lysis buffer (20 mM Tris-HCl (pH 7.4), 135 mM NaCl, 1.5 mM MgCl_2_, Triton 1%, Glycerol 10%, 1x Protease inhibitor (Roche)). Immunoblots were probed with antibodies against p53 (Santa Cruz sc126), Vinculin (Cambridge Bioscience 66305) and cyclin E (Abcam ab3927) and developed using goat antimouse (Cell Signalling 7076S) or goat anti-rabbit (Santa Cruz sc-2004) IgG HRP conjugated antibodies, using a Chemidoc (GE Healthcare).

### Flow cytometry

Cells were fixed in 4% formaldehyde for 7 min, permeabilised with 0.2% Triton X-100 for 2 min, stained with DAPI, and analysed using BD FACS Diva 8.2. RPE-1 cells were used to calibrate FACS analysis to generate a profile for DNA signal peaks, corresponding to a diploid cell line. RPE1-H2B-RFP and parental RPE-1 cells were then mixed and analysed together, for a direct comparison, to verify that H2B tagging did not alter the expected peaks. RPE-H2B-RFP cells were then mixed with known near-diploid or aneuploid cell lines (HCT116 and SW1116) to verify that the FACS analysis could distinguish that RFP positive cells gave the expected profile compared to the RFP negative cells when analysed together. RPE1-H2B-RFP cells were then mixed with individual HGSC cell lines, to see whether each HGSC cell line overlapped the diploid DNA signature or differed, indicating aneuploidy.

### Statistical Analysis

Statistical tests were carried out where indicated in figure legends, as either an unpaired t-test with Welch’s correction, or a one-way ANOVA with Tukey’s multiple comparison (where every cell line was compared to every other cell line; we only reported the results of pairwise comparisons between FNE1 and each individual HGSC cell line). Asterisks have been used to denote the significance value between experimental conditions adhering to the following nomenclature: p<0.05 (*); p<0.01 (**); p<0.001 (***); p<0.0001 (****). For non-significant differences, we report the actual p-value. All calculations were carried out using software (Graphpad Prism 7.0).

## Supplementary materials for Tamura et al, 2019

(one supplemental movie, two supplemental tables, five supplemental figures, see below) Additional materials:

**Movie S1: HGSC cells exhibit elevated microtubule assembly rates**. Shown is a EB3-GFP microtubule tip tracking movie from a COV318 HGSC cell treated with monastrol and filmed every 2 s (see Materials and methods for details).

**Table S2**: Mutations in *TP53, BRCA1* and *BRCA2* identified from whole genome sequencing of six HGSC cell lines (excel file)

**Table S1:**
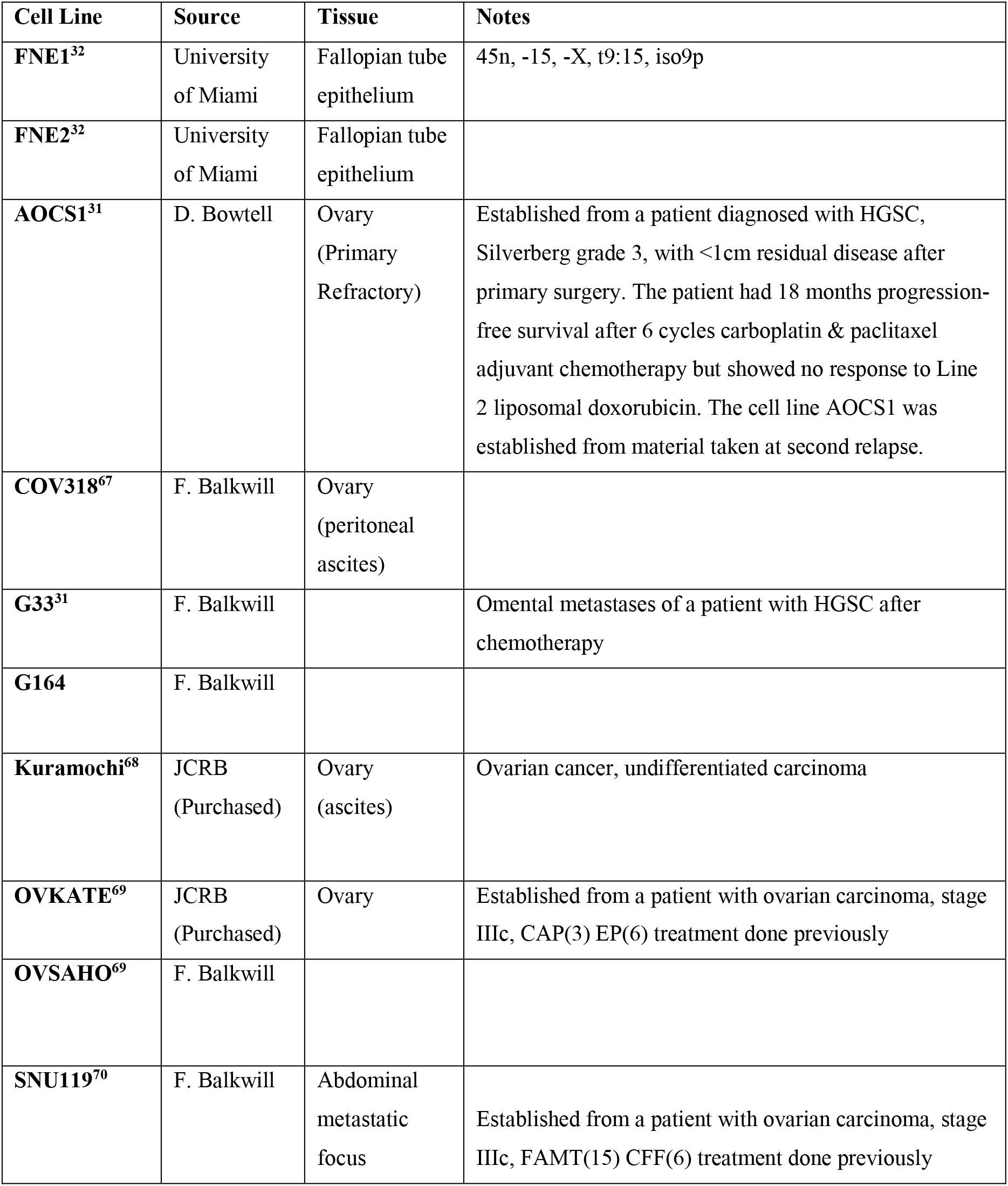
Details of HGSC cell lines used in this study.

**Figure S1 (relating to Figure 1):**
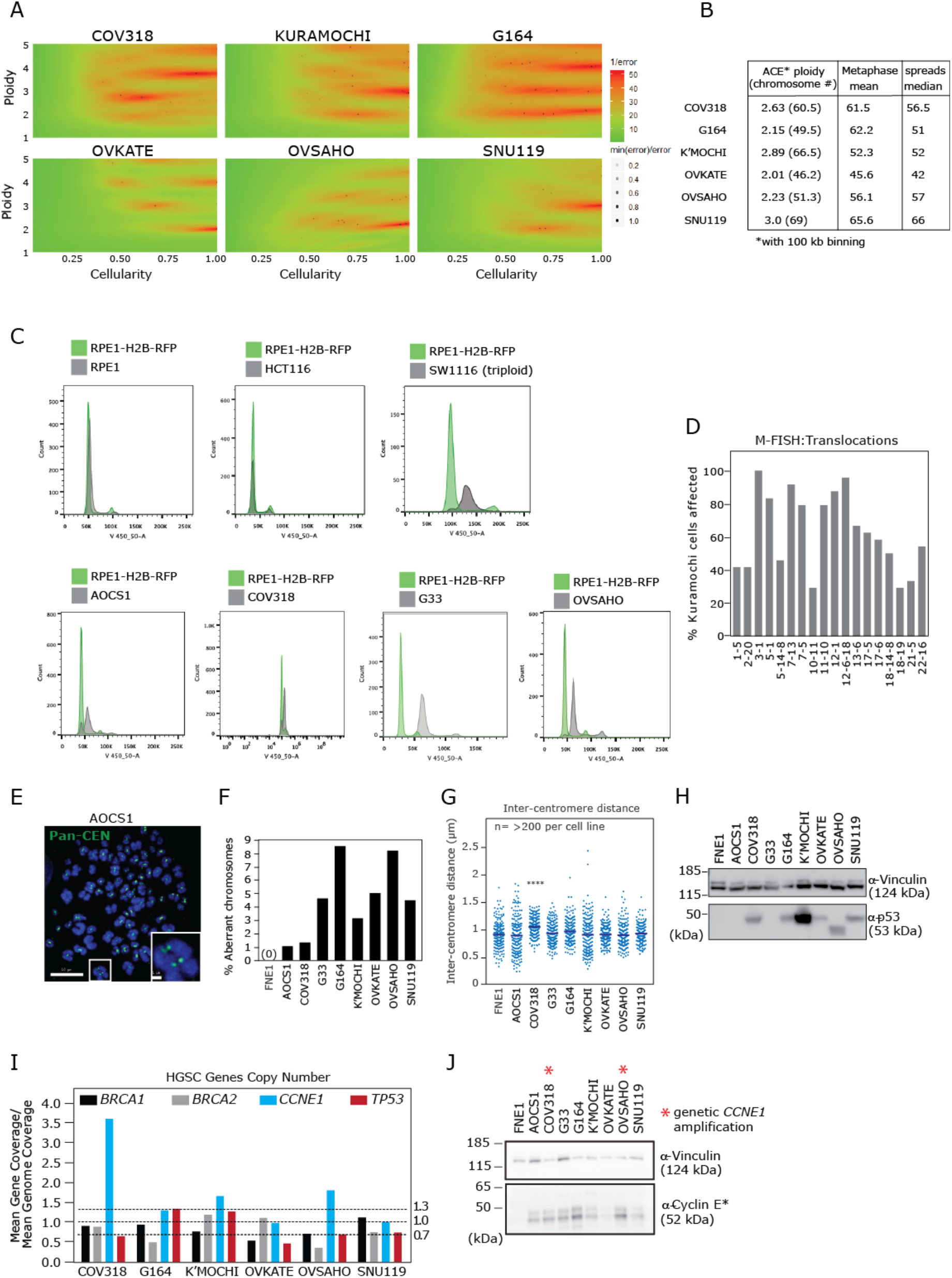
**A)** Matrix of errors plots derived from the loopsquaremodel function of ACE to indicate ploidy and cellularity probabilities (see Materials and methods for details). **B)** Table showing the ploidy values obtained from ACE analysis compared to mean and median chromosome counts from metaphase chromosome spreads. **C)** Flow cytometry analysis of HGSC compared to diploid cell lines. Each ‘test’ cell line sample was spiked with diploid RPE1-H2B-RFP cells to provide an internal control. CRC cell lines HCT116 (near diploid) and SW1116 (near triploid) were also analysed as positive controls for diploidy and triploidy respectively. **D)** Analysis of M-FISH images of 24 Kuramochi metaphase spreads (see **Figure 1**). Percentage of cells showing each indicated translocation. **E)** Example of a metaphase spread probed with pan-centromere FISH probes (Pan-CEN; green). Scale bar 10 μm. **F)** Percentage of all chromosomes analysed from metaphase spreads which demonstrated a structural defect (i.e. dicentric or acentric) (893-1511 chromosomes analysed across two experiments for each cell line). **G)** Intercentromere distance measured from metaphase spreads probed with pan-CEN FISH probes. **H)** Western Blot indicating protein levels of p53 in untreated cell lines as indicated. Vinculin used as loading control. **I)** Graph showing DNA copy number alterations of key genes derived from calculating gene reads coverage as a function of mean genome read depth (see Methods for details). **J)** Western Blot indicating expression levels of Cyclin E in untreated cell lines as indicated. Vinculin used as loading control. Asterisk indicates that Cyclin E is often present in tumour-specific lower molecular weight forms^71^.

**Figure S2 (relating to Figure 2):**
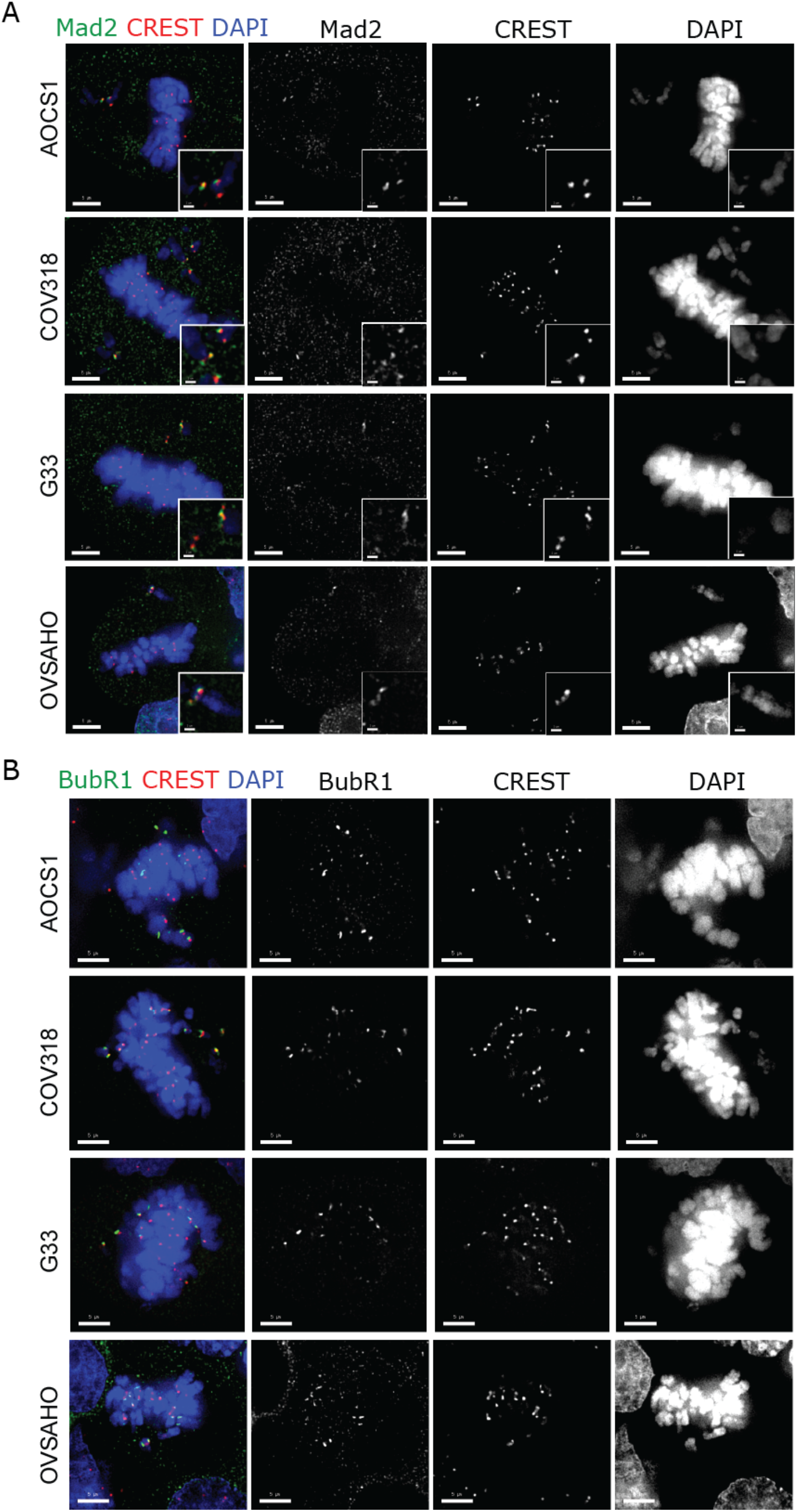
The spindle assembly checkpoint is functional in HGSC. **A,B)** Prometaphase cells from HGSC cell lines as indicated were probed with antibodies against CREST (red, marks centromeres) and either Mad2 (in A) or BubR1 (in B), components of the spindle assembly checkpoint that accumulate on unattached kinetochores and delay anaphase onset. Unaligned centric chromosomes demonstrated robust loading of Mad2 and BubR1.

**Figure S3 (relating to Figures 3 and 4):**
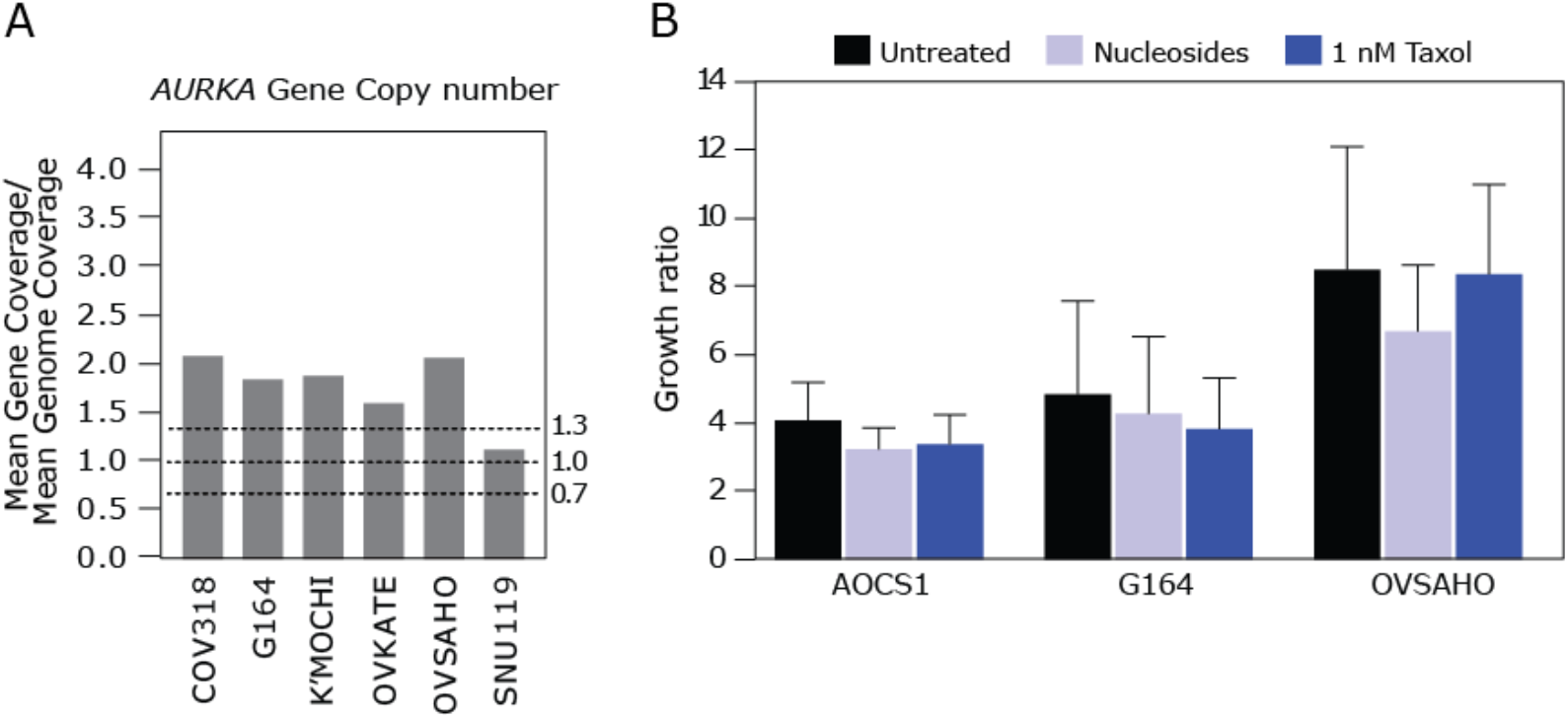
*AURKA* is amplified in HGSC, and long-term treatment with low dose taxol or nucleosides does not affect cell proliferation. **A)** Graph showing DNA copy number alterations of Aurora Kinase A (*AURKA*) derived from calculating gene reads coverage as a function of mean genome read depth (see Methods for details). **B)** Three HGSC cell lines were treated with taxol or nucleosides as indicated before measuring cell proliferation using Incucyte confluence measurement for one week. No significant differences were observed between growth rates of untreated and treated cells for each cell line.

**Figure S4 (relating to Figure 4):**
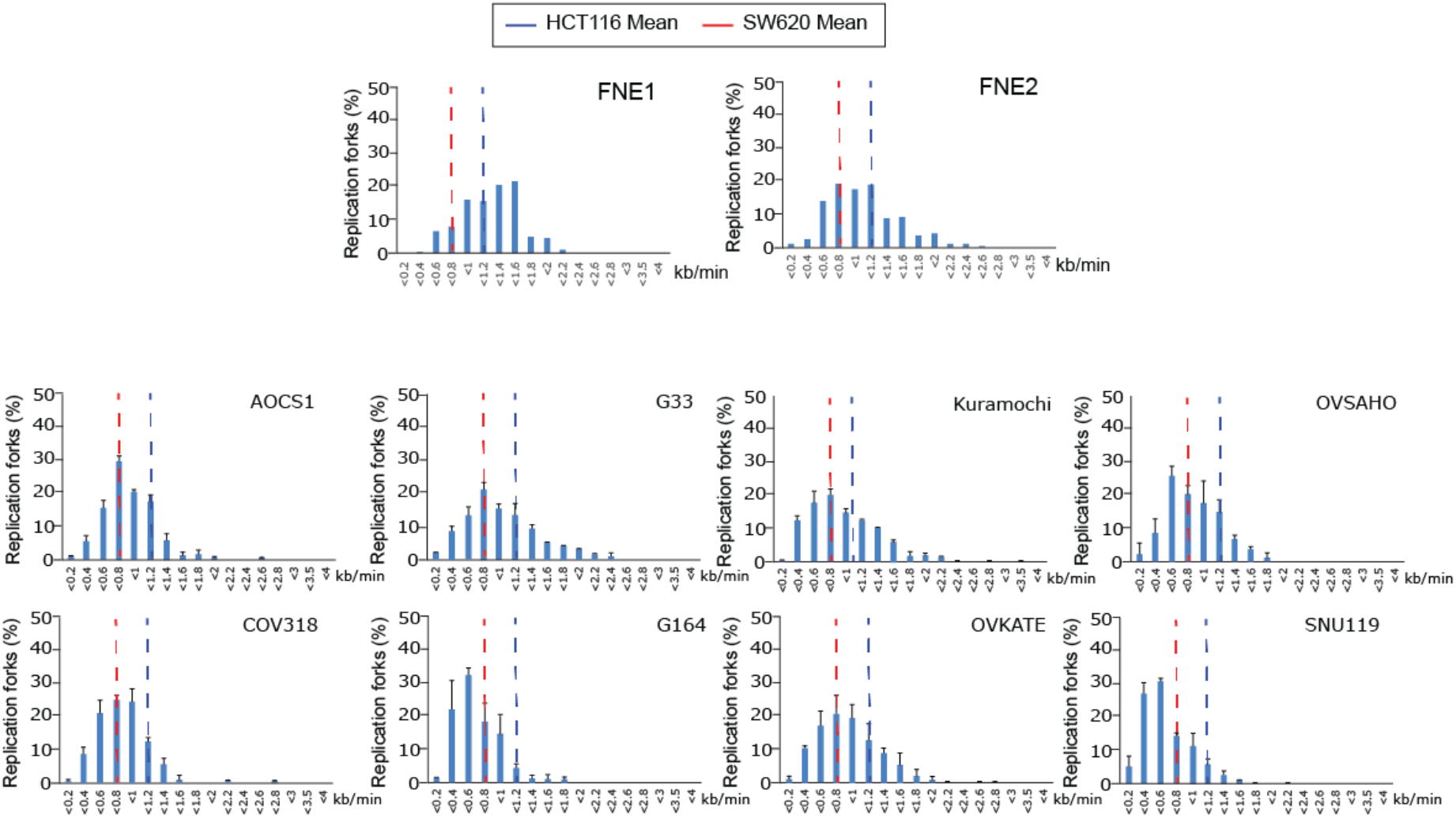
HGSC cell lines exhibit reduced replication fork rates. DNA fibre analysis was performed for HGSC cell lines as indicated. Single fibres were measured (99-202 fibres per cell line in total) and histograms showing the distribution of fork rates are shown. Also indicated at the mean fork rates of CRC lines HCT116 (CIN negative; blue dashed line) and SW620 (CIN positive; red dashed line).

**Figure S5 (relating to Discussion):**
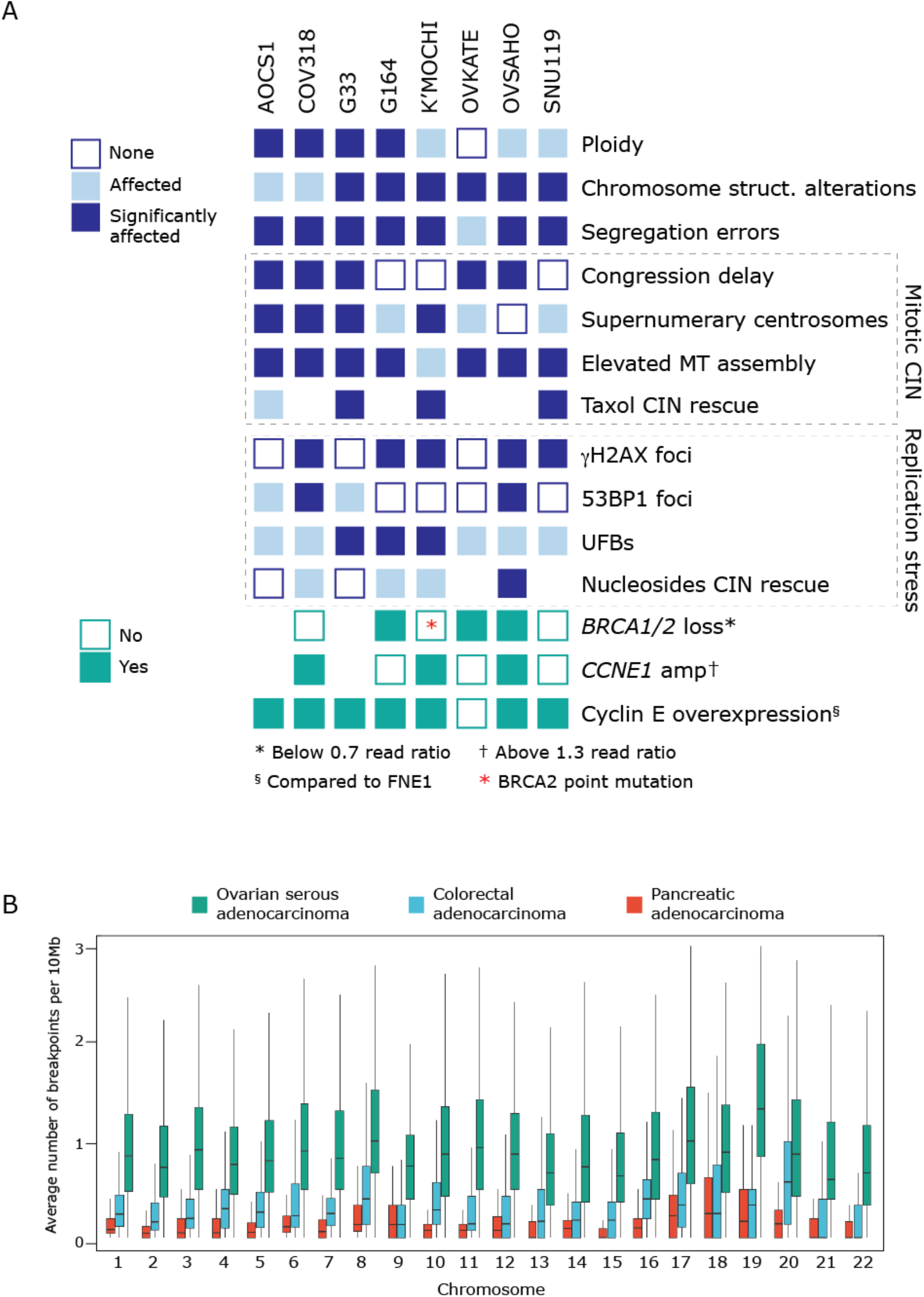
Phenotype summary and high rates of breakpoints in HGSC compared to other cancers. **A)** Diagrammatic summary of phenotypes across all cell lines. Chromosome structural (struct.) alterations refers to dicentric or acentric chromosomes. **B)** The number of breakpoints per 10 MB genome segments per chromosome, as calculated from the segmentation copy number data of the TCGA Ovarian Serous Cystadenocarcinoma, Colorectal Adenocarcinoma and Pancreatic Adenocarcinoma datasets (TCGA-OV, TCGA-COADREAD and TCGA-PAAD respectively). TCGA segmentation copy number data was downloaded from cBioportal^72,73^ and the number of breakpoints were calculated using an R-script adapted from available code as written by Macintyre et al^20^.

## Notes

The authors declare no potential conflicts of interest.

